# Time activity budget of White-rumped vulture and Slender-billed vulture during breeding in captivity

**DOI:** 10.64898/2025.12.09.693217

**Authors:** Sachin P. Ranade

## Abstract

Time activity budget is a widely used method to study the behavioral patterns of a range of animals including mammals and birds. The time budget of breeding birds offers valuable information regarding parental participation, incubation and chick rearing. The *Gyps* vultures were not intensively studied until recently. The Vulture Conservation Breeding Program of the Bombay Natural History Society is the first to hold and breed the resident *Gyps* vulture populations in captivity for their reintroduction in the wild. Hence, the White-rumped vulture and Slender-billed vulture were scientifically maintained in captivity and studied for their time activity budget. The aim of the study was understanding the role played by both sexes in breeding and derive the major activities of the parents and their offspring in the nest. The nesting vultures were observed with binoculars and through CCTV camera installed in the aviaries, during the day time from 06:00 to 17:30 hours to record activity of each parent at five-minute intervals. The activities for male and female individuals were divided into pre-hatching and post-hatching activities. Activities such as nest building, incubation, brooding and taking care of nestlings were recorded. Incubation and brooding were found to be the major activities of the breeding birds in both the *Gyps* species included in this study. Male and female individuals equally contributed for the incubation of the egg underlining their monomorphic nature.

## Introduction

Time activity budget method is used to study the behavior patterns of varied animal taxa from insects to mammals and birds. This method serves to derive information on various issues of an individual as well as the group. A few examples of use of activity budget in mammals are ─ comparison of food intake, rest and locomotion, among various age-sex groups in Howler monkey troop *Alouatta caraya* (Bicca-Marques and Calegaro-Marques 1994), study of foraging ecology and home range in Arunachal macaque *Macaca munzala* (Suresh Kumar *et al* 2007), time allocation in female Chimpanzees *Pan troglodytes schweinfurthii* during sexual cycle (Matsumoto-Oda & Oda 1998), travelling, feeding & socialization in Dolphin *Delphinus delphis* (Neumann 2001), and daily activity pattern in Spotted Hyena *Crocuta crocuta* (Kolowski 2007). The time budget of breeding birds offers valuable information about parental participation, incubation and chick rearing; few examples of such studies are Black-necked Cranes *Grus nigricollis* in China (Yang et al 2007), White-napped Cranes *Grus vipio* in Mongolia (Bradter et al. 2007), Gadwall *Anas strepera* (Dwyer 1975) and in the colonial sea-bird Common murre *Uria aalge* (Harding *et al*. 2007). The time budget of non-breeding birds reveals information about aspects like forging, locomotion and maintenance e.g. in Cape cormorant *Phalacrocorax capensis* (Ryan et al. 2010), in Teal *Anas crecca* (Khan *et al*. 1994), in the Tibetan eared pheasants *Crossoption harmani* (Lu & Zheng 2009) and in Helmeted guinea fowl *Numidia meleagris* (Kumssa & Bekele 2013). This method has been used to relate the time spent on grooming in different bird species and their parasite load (Cotgreave & Clayton 1994). The time budget study in some cases helps to understand the habitat use and anthropogenic pressure to the species as was studied in Lesser Spotted eagle *Aquila pomarina* (Meyburg *et al*. 2004). In conservation biology, it is used to evaluate reintroduction success of a species by comparing the pre-& post release activity pattern, for example this type of study was carried out in ‘Takhi’ *Equus ferus* in Mongolia (Boyd & Bandi 2002). The time budget and activity pattern monitoring also has economic application as it is used for understanding the needs & behavior of birds reared for meat/human consumption. Apart from its use in poultry industry, it has also been used in commercially raising of Ostriches *Struthio camelus* (Degen *et al* 1989).

‘Focal animal sampling’ is a classical method for collecting data required for the time budget and activity pattern study, but now-a-days use of satellite telemetry and data loggers are frequently used. The satellite telemetry has taken this study to the next level, for example in California condor, *Gymnogyps californianus,* automated satellite telemetry provided continuous position reporting and unbiased spatial coverage, along with thematic content such as the time, place and duration of particular activities (Cogan *et al*. 2012). Use of accelerometer and pop-up satellite tags (PSATs) are being developed to track the activity in aquatic habitat, e.g., in Sturgeon fish *Acipenser brevirostrum* (Boroell *et al*. 2016). Active acoustic tracking is also used to understand the movement and habitat usage as in case of the blue-spotted flathead fish, *Platycephalus caeruleopunctatus* (Fetterplace *et al* 2016).

The *Gyps* vultures were not intensively studied until the last decade of twentieth century. A few studies were carried out on the time-budget of the wild vulture population such as Griffon vulture, *Gyps fulvus* (Xirouchakis & Andreou 2009) and White-rumped vulture (Ahmad 2004). Owing to the drastic decline in resident *Gyps* vultures due to the veterinary use of diclofenac (Prakash 1999, Green *et al* 2004), the Bombay Natural History Society initiated the captive breeding program of *Gyps* vultures. The Vulture Conservation Breeding Program of the Bombay Natural History Society is the first to hold and breed the resident *Gyps* vulture populations in captivity for their reintroduction into the wild. Hence, the White-rumped vulture (Oriental White-backed vulture or White-backed vulture) and Slender-billed vulture were scientifically maintained in captivity and studied for their time activity budget. The study was planned in a way to provide information about the *Gyps* vultures facilitating understanding of their behavior which would be helpful in the management practices for the Vulture Conservation Breeding Program. The observations were recorded at the colony aviary at the Vulture Conservation Breeding Center, Rajabhatkhawa, West Bengal during the breeding season 2010-2011.

The key objectives of the study were:

1. Understanding the role played by both sexes in breeding.
2. Find out the major activities of the parents and their offspring in the nest.

## Material and Methods

The time budget and activity pattern of the breeding vultures were studied in captivity for both the species ─ White-rumped vulture, *Gyps bengalensis* and slender-billed vulture, *Gyps tenuirostris*. The flock of captive wild vultures was observed for their breeding activities in the colony aviary. The colony aviary is a huge aviary (100 ft long, 40 ft wide and 20 ft high) which provided a habitat which was close to natural conditions. The nesting vultures were observed through one sided glass windows with the help of binoculars and through the monitor of CCTV camera installed in the aviaries, during the day time from 06:00 to 17:30 hours. The instantaneous scan method was followed (Altmann, 1974, Palmer et al. 2001) to sample activity of each parent at five-minute intervals. The activities were divided into pre-hatching and post-hatching activities, which were recorded during three time periods during the day e.g. morning 6:00 to 10:00 hours, mid-day 10:00 to 14:00 hours and evening 14:00 to 17:30 hours. The various activities including incubation, feeding, brooding, taking care of young ones and so on, were recorded. The time spent on various activities was compared during different hours of the day and time budget was generated.

The various activities noted in the adult vultures were as follows:

- Maintenance activities such as preening, bathing, scratching, sunning and wing exercise.
- Resting activities e.g. standing, standing on one leg, sleeping, squatting, resting in the nest, resting while sitting on hocks.
- Feeding activities like food search, actual feeding, regurgitating in the nest and feeding the chick.
- Nest building such as nest material collection, nest material arranging.
- Mating of the pair.
- Incubation of egg.
- Activities of taking care of the nestlings such as brooding and feeding the nestlings.

## Results

The White-rumped vulture was observed for 1797 hours while the Slender-billed vulture was observed for 996 hours during the breeding season 2010-11. As the vultures are not sexually dimorphic, male and female individuals were identified based on their courtship activities and ring numbers. The activities of adult birds were classified into pre-hatching and post-hatching. In case of post hatching, additional activities were observed – adults feeding the nestling and activities of the nestling.

### Activity pattern of breeding White-rumped vultures

Three nests of White-rumped vulture (N00, N26 and N28) were observed during this study. The activities of male, female and nestlings were recorded. The adult birds were absent from the nest for on an average 40% of the time under observation. It could be due to the bird being at far perch, or on ground or out of the view from its nest site. The time budget and activity pattern for various activities recorded in male and female White-rumped vultures is presented in Figures 1-12 and Tables 1-12.

**Table 1.** The activities of male White-rumped vulture (Nest N00) during the pre-hatching period. Total observations of 8970 minutes.

|  | Percentage time spent (Is that average of many observations) |  |  |
| --- | --- | --- | --- |
| Activity | 6 to 10 hours | 10 to 14 hours | 14 to 17.30 hours |
| Incubation | 56.7 | 71.7 | 51.4 |
| Perch on nest | 6.3 | 7.6 | 7.9 |
| Perch | 34.4 | 16.8 | 35.5 |
| Perch on ground | 1.2 | 2.6 | 0.6 |
| Feeding | 0 | 0.4 | 3 |
| Bath | 0 | 0 | 0 |
| Nest material collection | 0.4 | 0.1 | 0.4 |
| Nest arrangement | 0.8 | 0.9 | 1.1 |
| Copulation | 0.2 | 0 | 0.2 |

**Figure 1.**
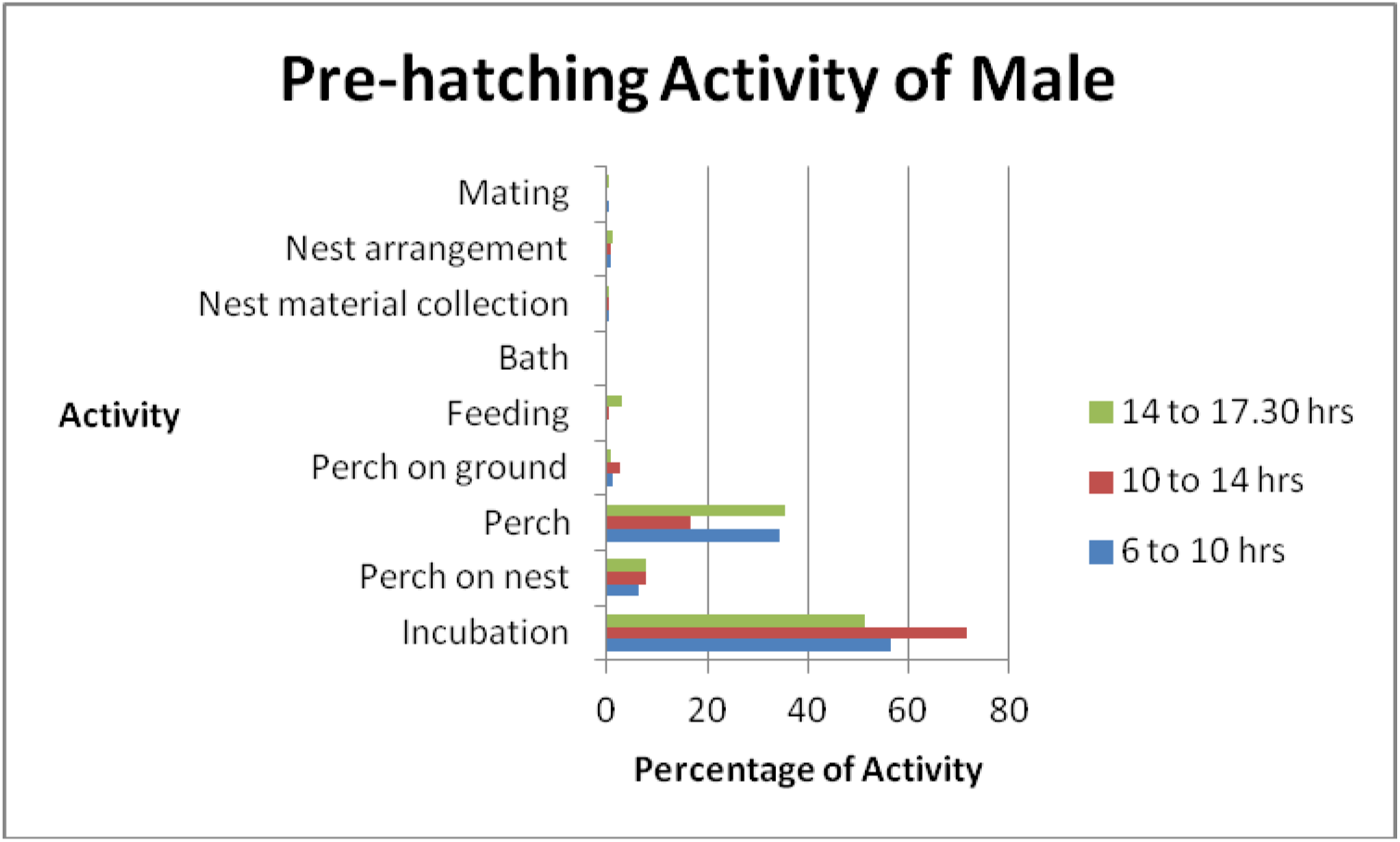
The activities of male White-rumped vulture (Nest N00) during the pre-hatching period.

**Table 2.** The activities of female White-rumped vulture (Nest N00) during the pre-hatching period. Total observations of 9545 minutes.

| Activity | Percentage activity |  |  |
| --- | --- | --- | --- |
|  | 6 to 10 hours | 10 to 14 hours | 14 to 17.30 hours |
| Incubation | 74.5 | 76.2 | 80.8 |
| Perch on nest | 5.7 | 3.5 | 3.5 |
| Perch | 15 | 11.6 | 6.8 |
| Perch on ground | 2.2 | 5 | 0.9 |
| Feeding | 1.2 | 2.7 | 6.5 |
| Bath | 0 | 0.3 | 0 |
| Nest material collection | 0 | 0.1 | 0 |
| Nest arrangement | 1.2 | 0.7 | 1.4 |
| Mating | 0.2 | 0 | 0.2 |

**Figure 2.**
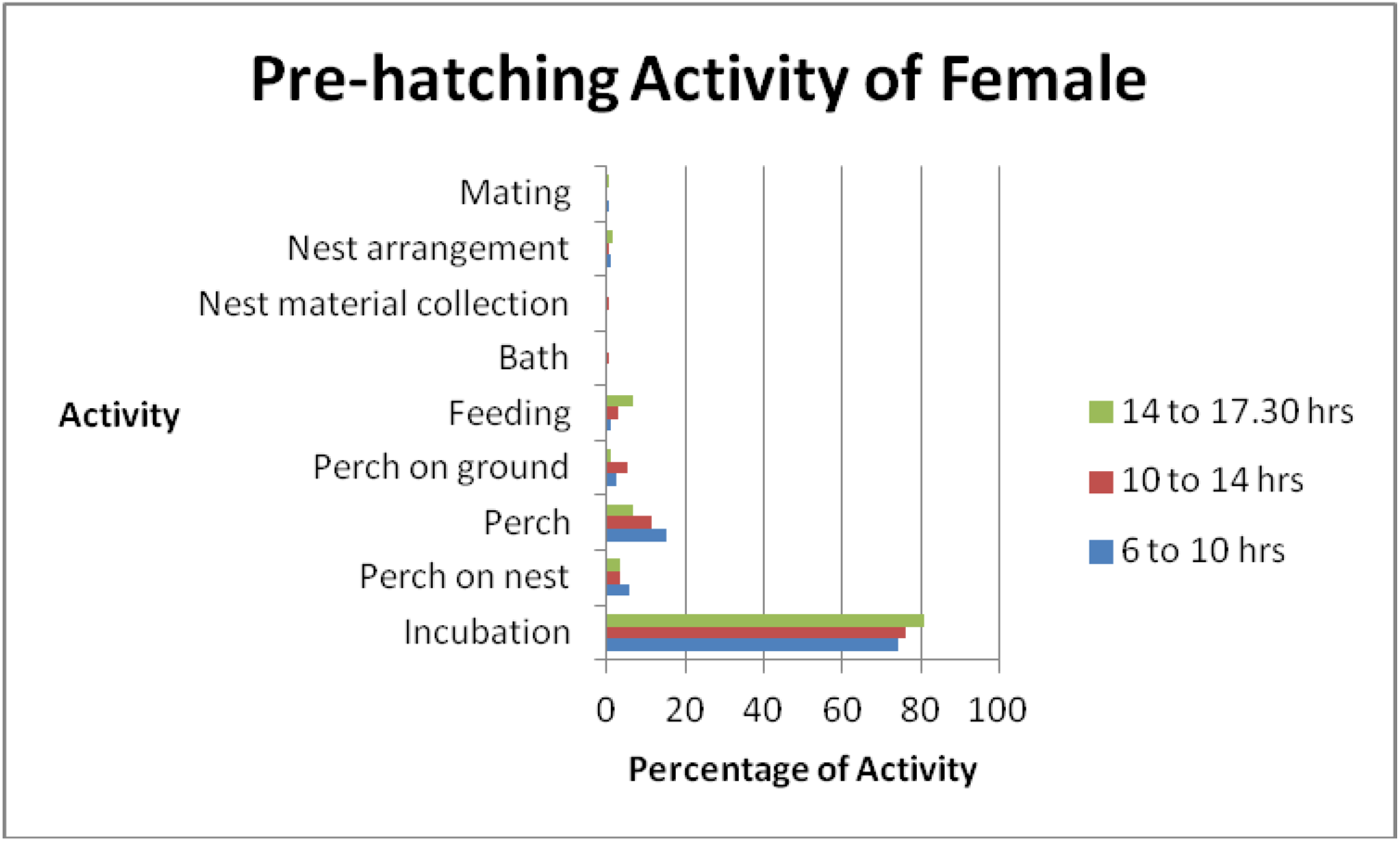
The activities of female White-rumped vulture (Nest N00) during the pre-hatching period.

**Table 3.** The activities of male White-rumped vulture (Nest N00) during the post-hatching period. Total observations of 3110 minutes.

| Activity | Percentage activity |  |  |
| --- | --- | --- | --- |
|  | 6 to 10 hours | 10 to 14 hours | 14 to 17.30 hours |
| Brood | 24.2 | 9.8 | 23.7 |
| Perch on nest | 15.8 | 15.6 | 25 |
| Perch | 50.5 | 62.3 | 43.6 |
| Perch on ground | 0 | 0 | 0 |
| Feeding | 2.1 | 0.4 | 0 |
| Bath | 0 | 0 | 0 |
| Nest material collection | 0 | 0 | 0 |
| Nest arrangement | 1.1 | 1.4 | 0 |
| Mating | 0 | 0 | 0 |
| Feeding the young | 4.2 | 10.1 | 7.7 |
| Preening the young | 2.1 | 0.4 | 0 |

**Figure 3.**
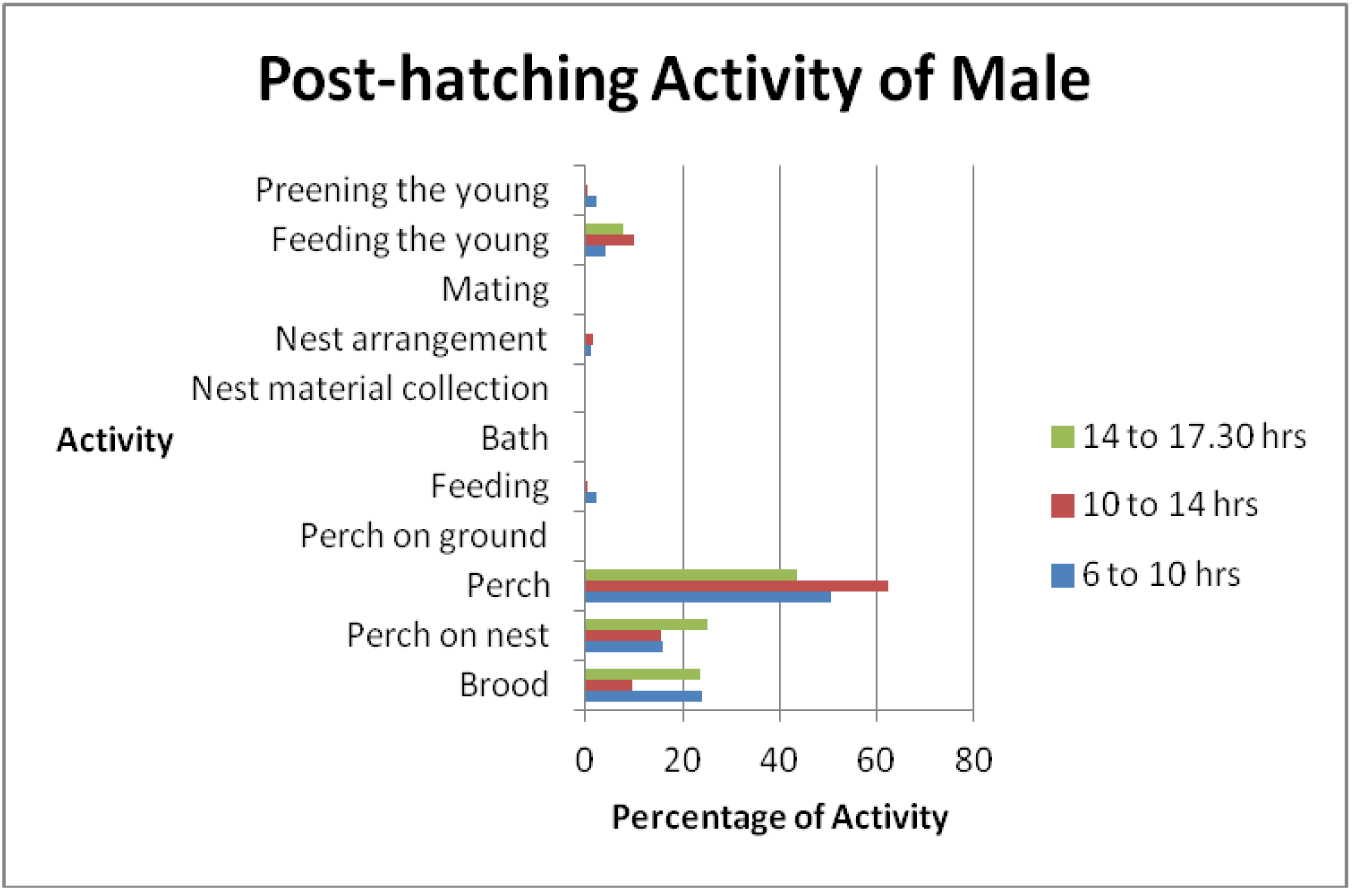
The activities of male White-rumped vulture (Nest N00) during the post-hatching period.

**Table 4.** The activity pattern female White-rumped vulture (Nest N00) during the post-hatching period. Total observations of 3005 minutes.

| Activity | Percentage activity |  |  |
| --- | --- | --- | --- |
|  | 6 to 10 hours | 10 to 14 hours | 14 to 17.30 hours |
| Brood | 26.5 | 32.7 | 51.9 |
| Perch on nest | 39.2 | 31.2 | 17.5 |
| Perch | 29.3 | 31.2 | 17.5 |
| Perch on ground | 0 | 0 | 0 |
| Feeding | 0 | 0 | 0 |
| Bath | 0 | 0 | 0 |
| Nest material collection | 0 | 0 | 0 |
| Nest arrangement | 0.6 | 0.8 | 1.3 |
| Mating | 0 | 0 | 0 |
| Feeding the young | 3.9 | 4.2 | 11.9 |
| Preening the young | 0.6 | 0 | 0 |

**Figure 4.**
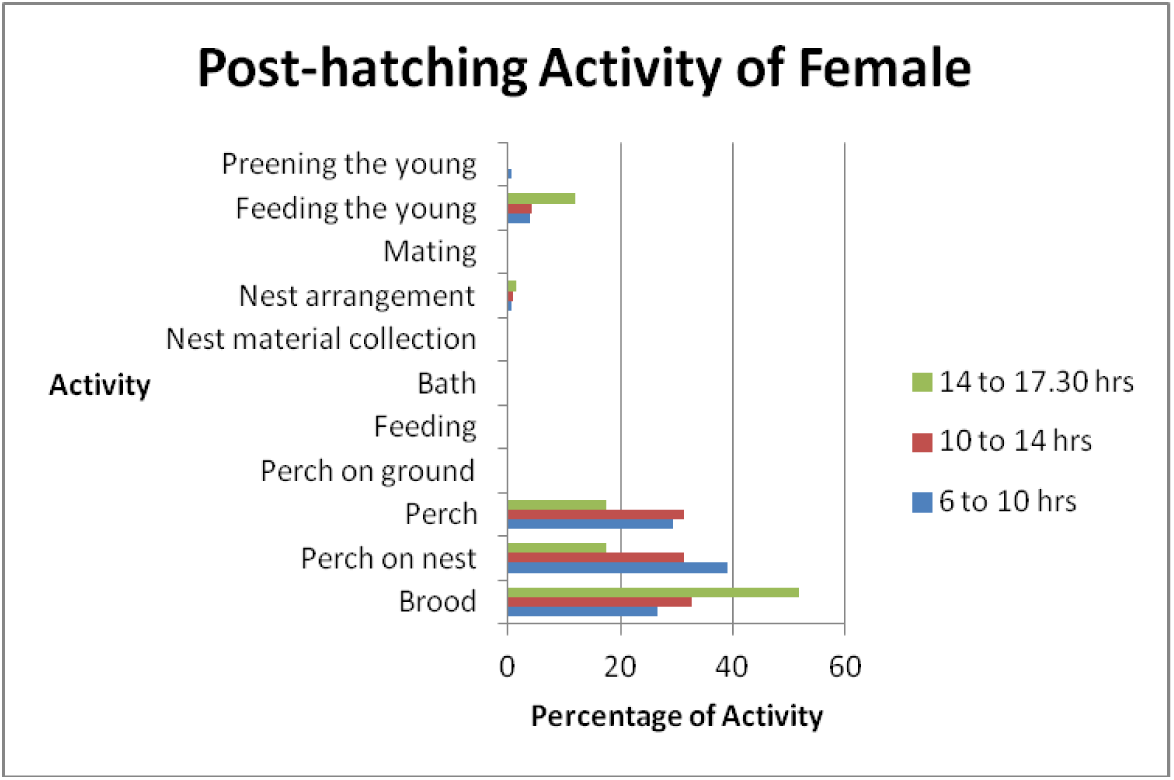
The activity pattern female White-rumped vulture (Nest N00) during the post-hatching period.

**Table 5.** The activities of male White-rumped vulture (Nest N26) during the pre-hatching period. Total observations of 12550 minutes.

| Activity | Percentage activity |  |  |
| --- | --- | --- | --- |
|  | 6 to 10<br>hours | 10 to 14<br>hours | 14 to 17.30<br>hours |
| Incubation | 71.5 | 64.5 | 61.7 |
| Perch on nest | 8.6 | 8 | 10.9 |
| Perch | 12.3 | 15.9 | 13.1 |
| Perch on ground | 2.7 | 8.4 | 4.1 |
| Feeding | 2.6 | 0.8 | 8.9 |
| Bath | 0 | 0.2 | 0 |
| Nest material<br>collection | 0.4 | 0.4 | 0.3 |
| Nest arrangement | 1 | 0.6 | 0.5 |
| Mating | 0.8 | 1.2 | 0.5 |

**Figure 5.**
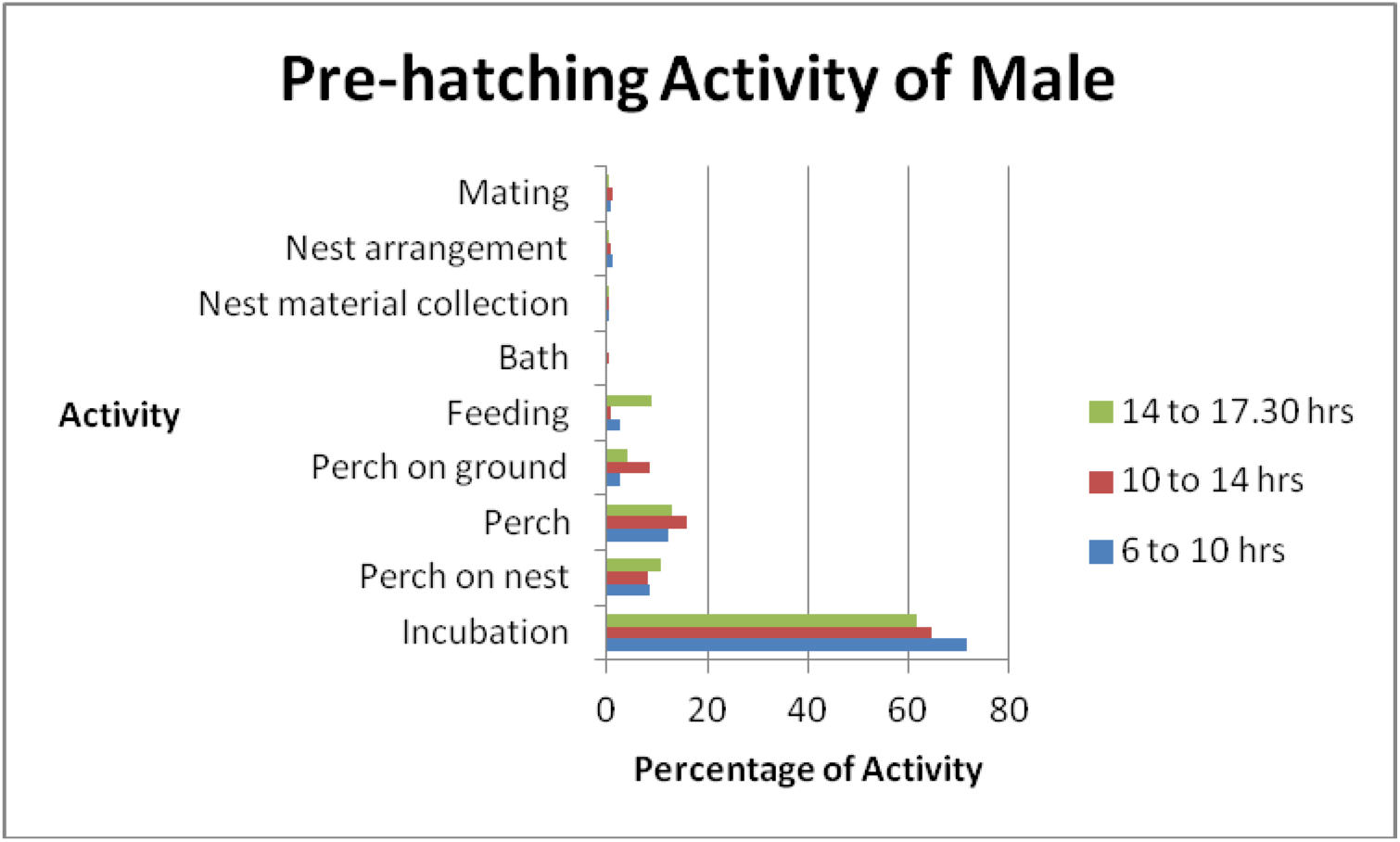
The activities of male White-rumped vulture (Nest N26) during the pre-hatching period.

**Table 6.**
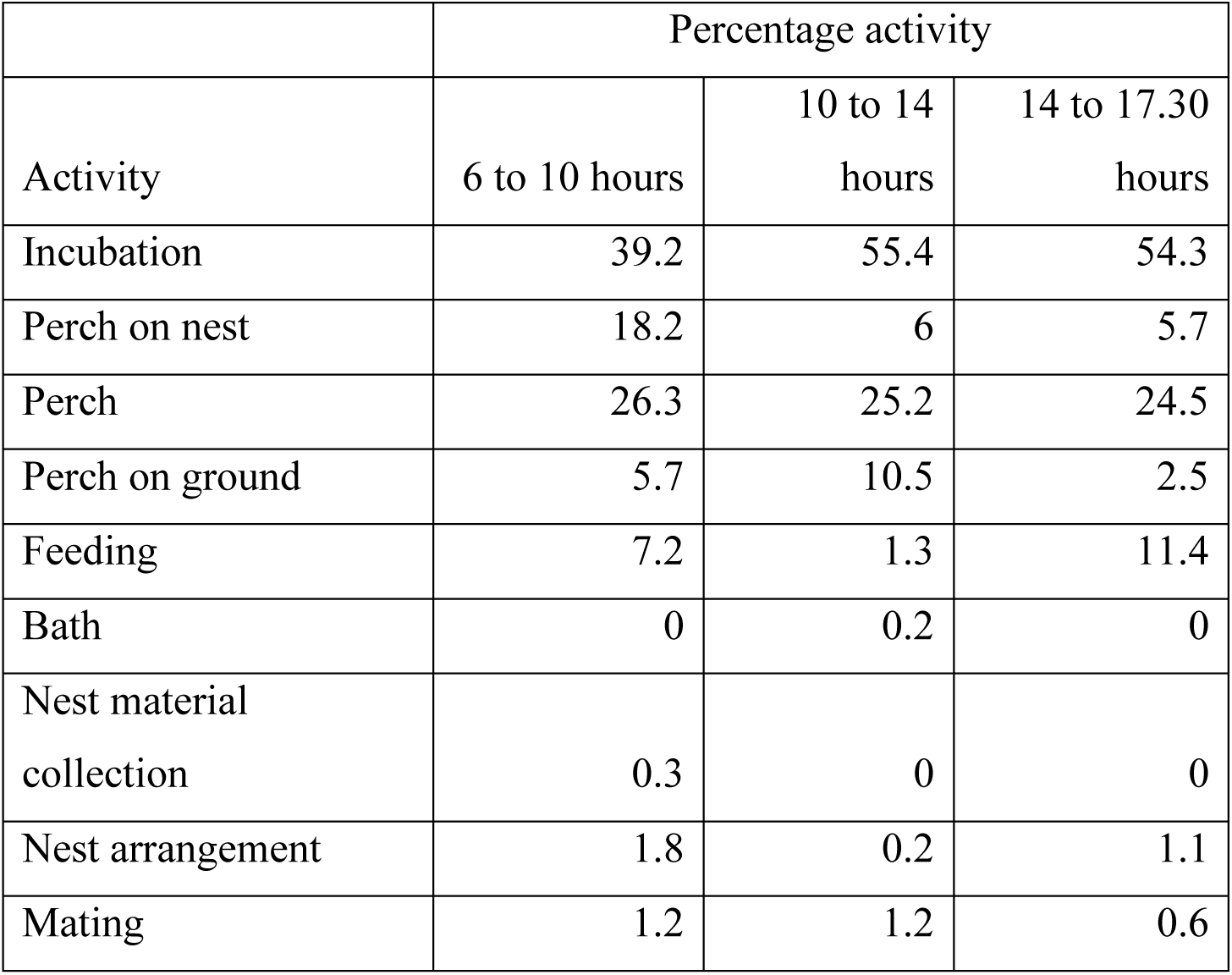
The activities of female White-rumped vulture (Nest N26) during the pre-hatching period. Total observations of 10610 minutes.

| Activity | Percentage activity |  |  |
| --- | --- | --- | --- |
|  | 6 to 10 hours | 10 to 14 hours | 14 to 17.30 hours |
| Incubation | 39.2 | 55.4 | 54.3 |
| Perch on nest | 18.2 | 6 | 5.7 |
| Perch | 26.3 | 25.2 | 24.5 |
| Perch on ground | 5.7 | 10.5 | 2.5 |
| Feeding | 7.2 | 1.3 | 11.4 |
| Bath | 0 | 0.2 | 0 |
| Nest material collection | 0.3 | 0 | 0 |
| Nest arrangement | 1.8 | 0.2 | 1.1 |
| Mating | 1.2 | 1.2 | 0.6 |

**Figure 6.**
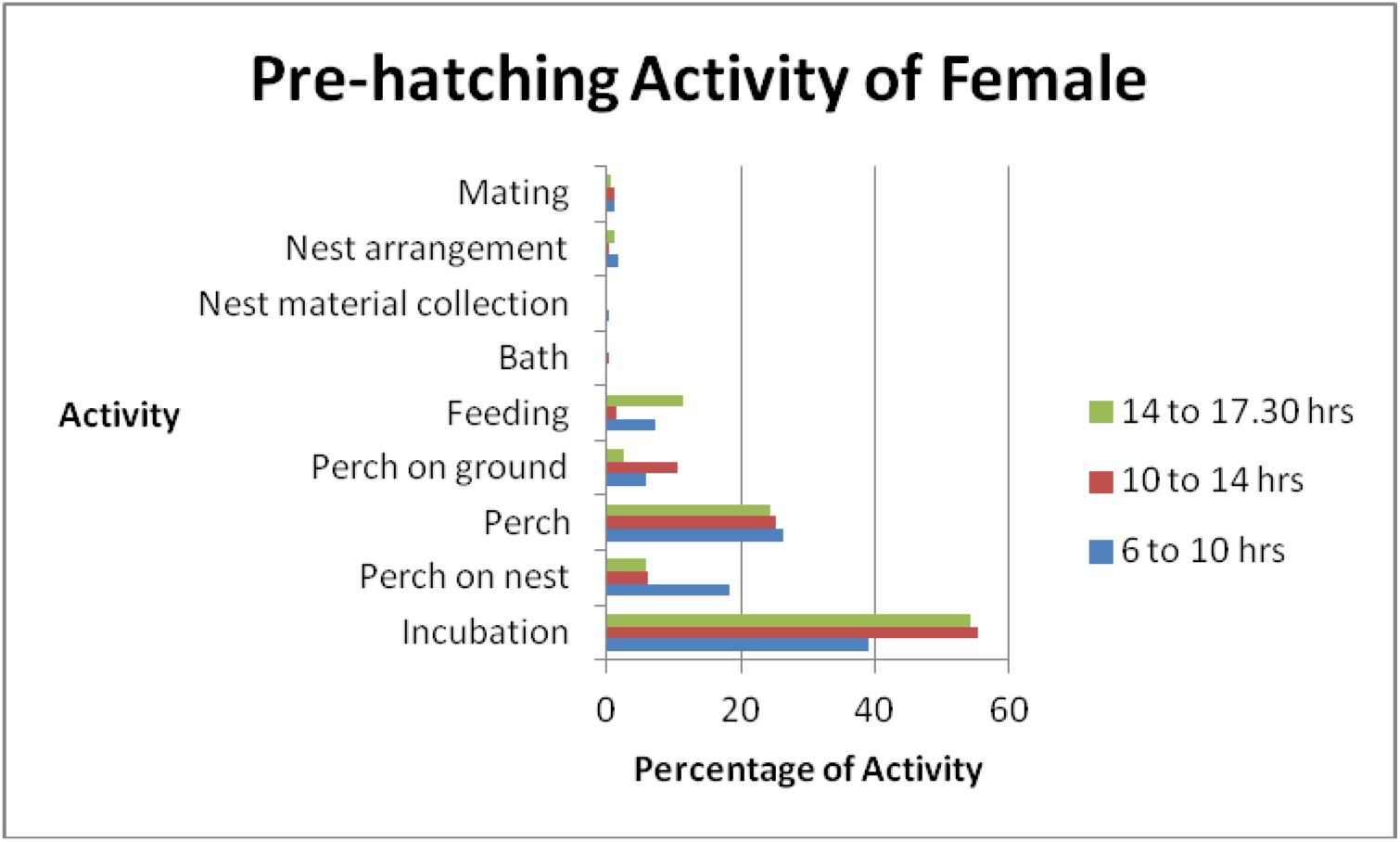
The activities of female White-rumped vulture (Nest N26) during the pre-hatching period.

**Table 7.** The activities of male White-rumped vulture (Nest N26) during the post-hatching period. Total observations of 5245 minutes.

| Activity | Percentage activity |  |  |
| --- | --- | --- | --- |
|  | 6 to 10 hours | 10 to 14 hours | 14 to 17.30 hours |
| Brood | 56.2 | 43.2 | 33.3 |
| Perch on nest | 21.9 | 36.9 | 33 |
| Perch | 8 | 6.6 | 17.2 |
| Perch on ground | 3.1 | 1.1 | 0 |
| Feeding | 0 | 0 | 0 |
| Bath | 0 | 0 | 0 |
| Nest material collection | 0 | 0 | 0 |
| Nest arrangement | 0.6 | 1.3 | 2.6 |
| Mating | 0.6 | 0 | 0.4 |
| Feeding the young | 9.3 | 10.3 | 13.1 |
| Preening the young | 0.3 | 0.7 | 0.4 |

**Figure 7.**
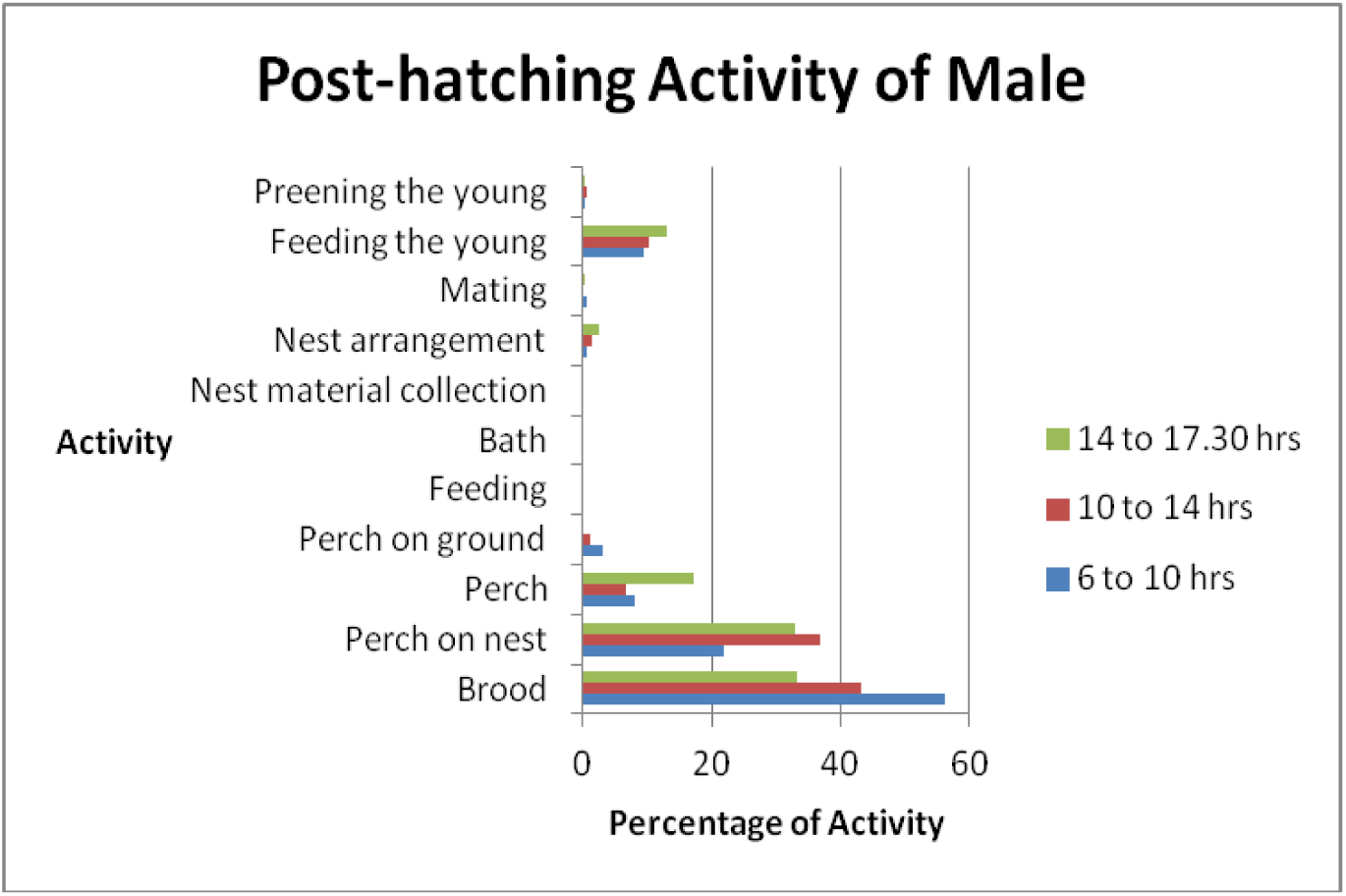
The activities of male White-rumped vulture (Nest N26) during the post-hatching period.

**Table 8.** The activities of female White-rumped vulture (Nest N26) during the post-hatching period. Total observations of 4480 minutes.

| Activity | Percentage activity |  |  |
| --- | --- | --- | --- |
|  | 6 to 10 hours | 10 to 14 hours | 14 to 17.30 hours |
| Brood | 14.7 | 14.1 | 21.1 |
| Perch on nest | 33.9 | 52.1 | 47.3 |
| Perch | 41.4 | 15.2 | 22.3 |
| Perch on ground | 0.4 | 7.6 | 0 |
| Feeding | 2 | 4.8 | 0 |
| Bath | 0 | 0 | 0 |
| Nest material collection | 0 | 0 | 0 |
| Nest arrangement | 1.6 | 0.7 | 0.6 |
| Mating | 0.8 | 0 | 0.3 |
| Feeding the young | 4 | 5.5 | 7.6 |
| Preening the young | 1.2 | 0 | 0.8 |

**Figure 8.**
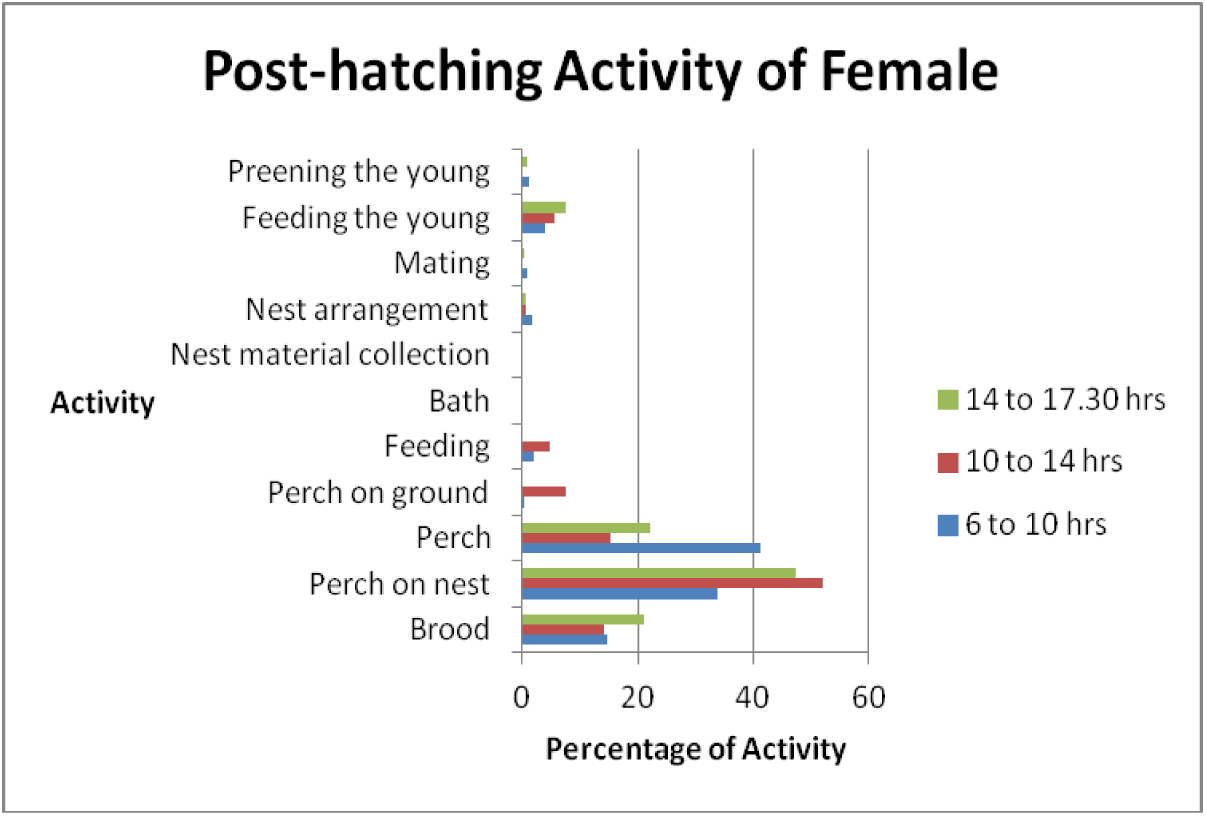
The activities of female White-rumped vulture (Nest N26) during the post-hatching period.

**Table 9.** The activities of male White-rumped vulture (Nest N28) during the pre-hatching period. Total observations of 13335 minutes.

| Activity | Percentage activity |  |  |
| --- | --- | --- | --- |
|  | 6 to 10<br>hours | 10 to 14<br>hours | 14 to 17.30<br>hours |
| Incubation | 59.3 | 65.3 | 53.5 |
| Perch on nest | 9.7 | 5.7 | 6.4 |
| Perch | 21.8 | 19.6 | 24.9 |
| Perch on ground | 4.8 | 6.9 | 4.8 |
| Feeding | 1.5 | 1.5 | 9.4 |
| Bath | 0 | 0.1 | 0 |
| Nest material collection | 1 | 0.3 | 0.2 |
| Nest arrangement | 1.4 | 0.4 | 0.6 |
| Mating | 0.6 | 0.3 | 0.2 |

**Figure 9.**
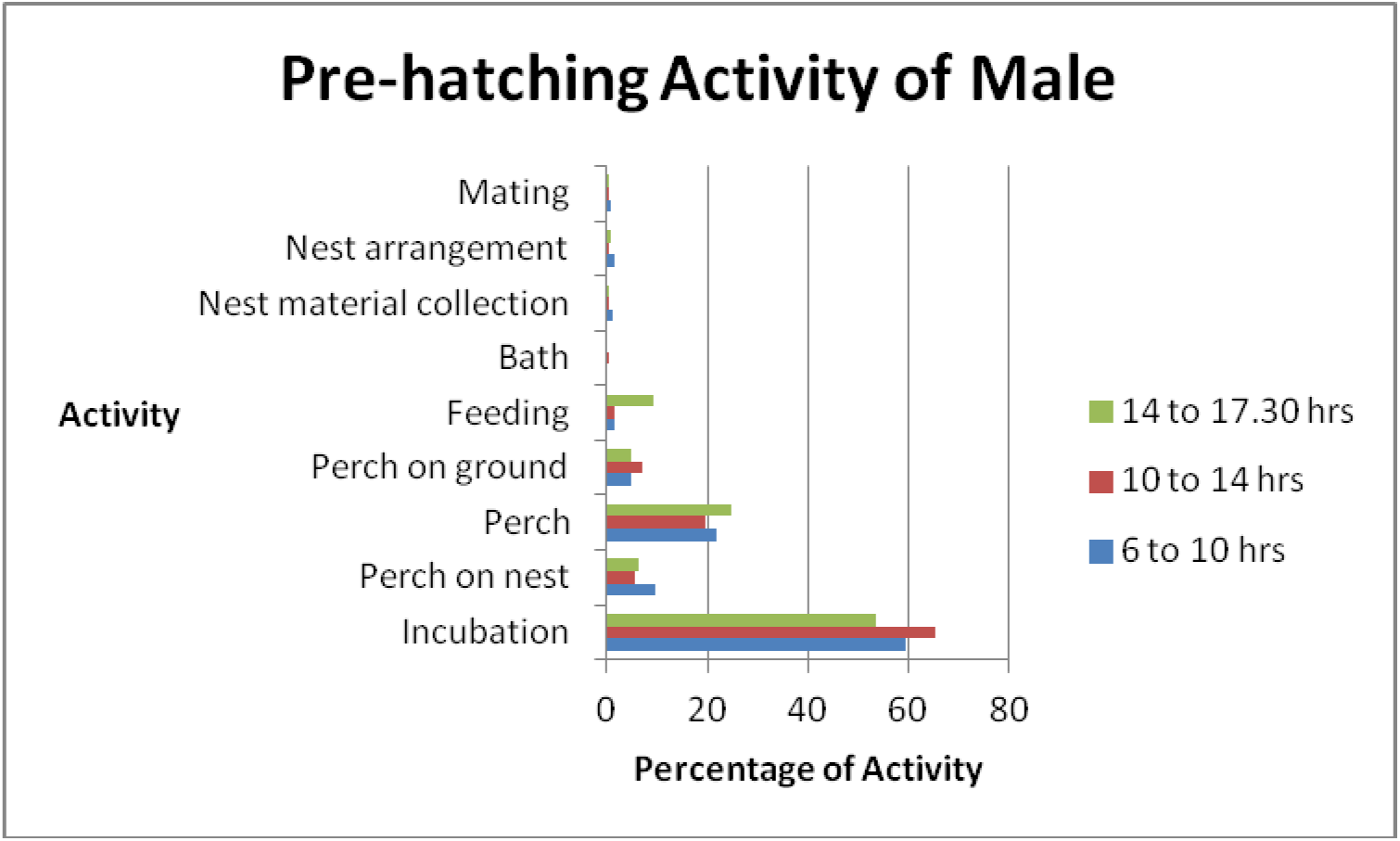
The activities of male White-rumped vulture (Nest N28) during the pre-hatching period.

**Table 10.** The activities of female White-rumped vulture (Nest N28) during the pre-hatching period. Total observations of 11525 minutes.

| Activity | Percentage activity |  |  |
| --- | --- | --- | --- |
|  | 6 to 10 hours | 10 to 14 hours | 14 to 17.30 hours |
| Incubation | 55.4 | 61.4 | 65.8 |
| Perch on nest | 20 | 7.4 | 8.3 |
| Perch | 13.7 | 20.1 | 20.2 |
| Perch on ground | 3.2 | 7.8 | 0.2 |
| Feeding | 3.9 | 2.2 | 4.6 |
| Bath | 0 | 0.2 | 0 |
| Nest material collection | 0.1 | 0 | 0 |
| Nest arrangement | 2.9 | 0.6 | 0.8 |
| Mating | 0.8 | 0.3 | 0.2 |

**Figure 10.**
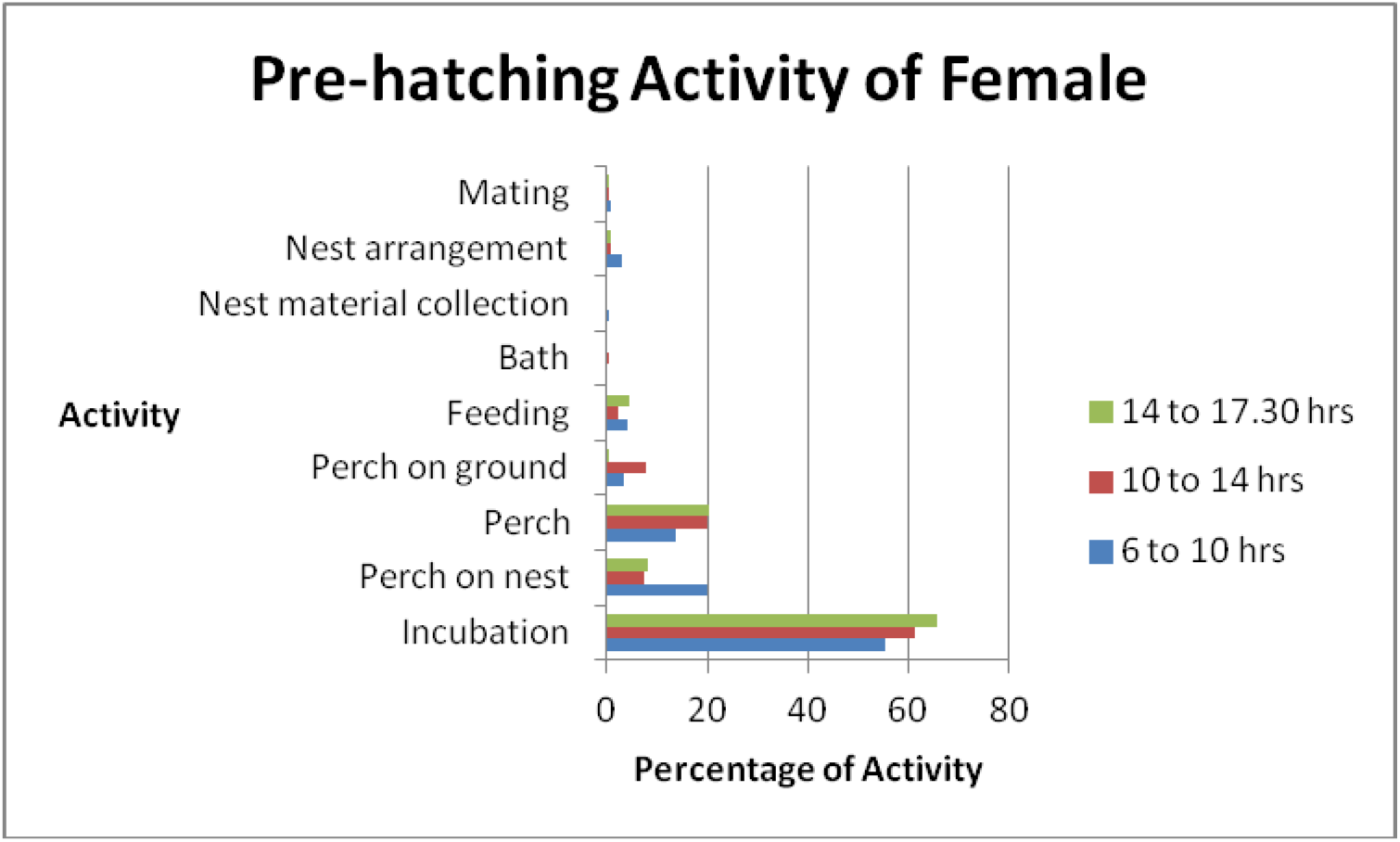
The activities of female White-rumped vulture (Nest N28) during the pre-hatching period.

**Table 11.** The activities of male White-rumped vulture (Nest N28) during the post-hatching period. Total observations of 4940 minutes.

| Activity | Percentage activity |  |  |
| --- | --- | --- | --- |
|  | 6 to 10 hours | 10 to 14 hours | 14 to 17.30 hours |
| Brood | 52.6 | 30.2 | 17.2 |
| Perch on nest | 17.1 | 28.6 | 54.8 |
| Perch | 23.1 | 30.4 | 19.4 |
| Perch on ground | 0.3 | 0 | 0 |
| Feeding | 0.6 | 0 | 0 |
| Bath | 0 | 0 | 0 |
| Nest material collection | 0.3 | 0.3 | 0 |
| Nest arrangement | 0.6 | 0.5 | 0.4 |
| Mating | 0 | 0 | 0 |
| Feeding the young | 5.3 | 10.1 | 8.2 |
| Preening the young | 0 | 0 | 0 |

**Figure 11.**
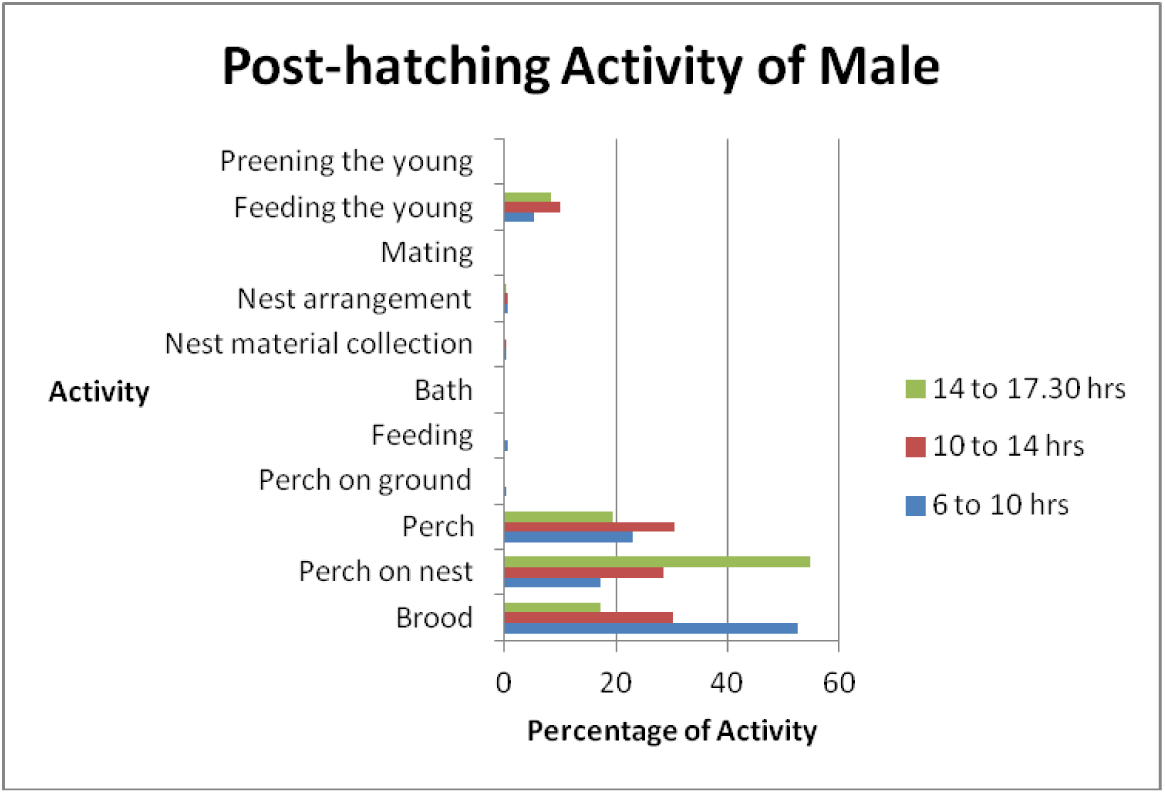
The activities of male White-rumped vulture (Nest N28) during the post-hatching period.

**Table 12.** The activities of female White-rumped vulture (Nest N28) during the post-hatching period. Total observations of 4680 minutes.

| Activity | Percentage activity |  |  |
| --- | --- | --- | --- |
|  | 6 to 10 hours | 10 to 14 hours | 14 to 17.30 hours |
| Brood | 28.6 | 26.1 | 18.9 |
| Perch on nest | 34.3 | 40.9 | 39.6 |
| Perch | 24.3 | 20.8 | 33.1 |
| Perch on ground | 1.4 | 0 | 0 |
| Feeding | 1.1 | 0 | 0 |
| Bath | 0 | 0 | 0 |
| Nest material collection | 0 | 0 | 0 |
| Nest arrangement | 1.1 | 2.2 | 1.2 |
| Mating | 0 | 0 | 0 |
| Feeding the young | 8.9 | 9.7 | 7.1 |
| Preening the young | 0.4 | 0.3 | 0 |

**Figure 12.**
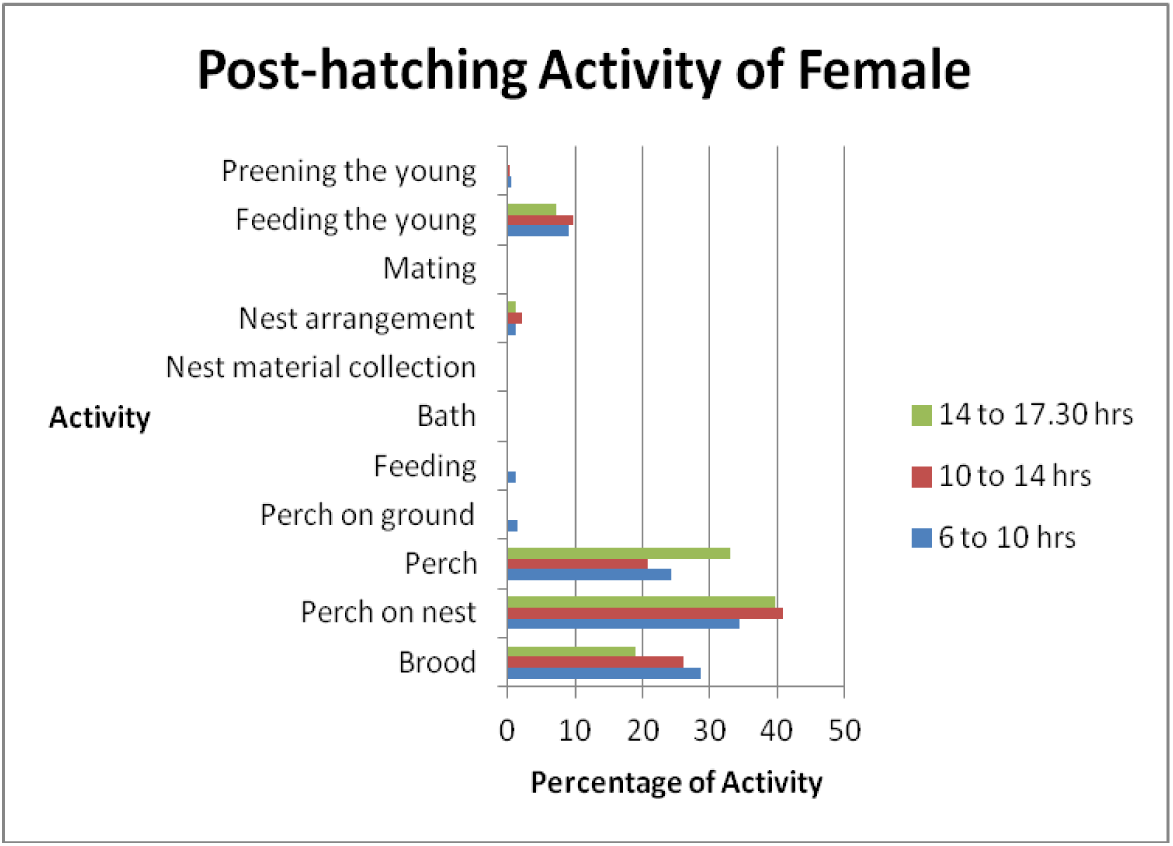
The activities of female White-rumped vulture (Nest N28) during the post-hatching period.

### Activity pattern of breeding Slender-billed vultures

Two nests of Slender-billed vulture N7 and N16, were monitored for the activity pattern in captivity. Total observations recorded were for 996 hours. The nests were constructed by the species in the topmost rows of provided nest ledges. During the observations, the nesting attempt was not successful. As there was the failure in hatching, the only pre-hatching activities are considered for both nests and both sexes. The time budget and activity pattern for various activities recorded in male and female Slender-billed vultures is presented in Figures 13- and Tables 13-.

**Table 13.**
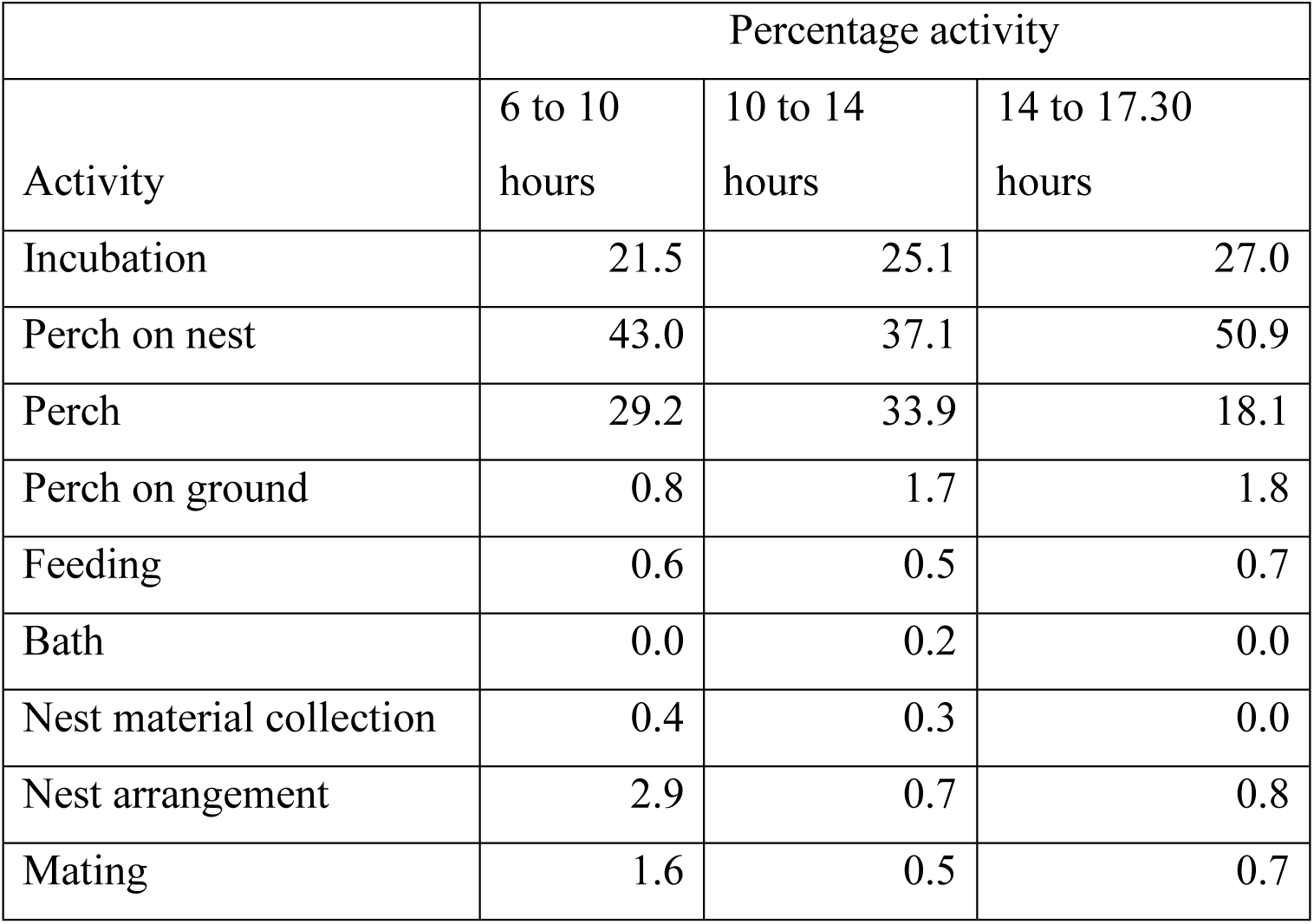
The activities of male Slender-billed vulture (Nest N7) during the pre-hatching period. Total observations of 20170 minutes.

| Activity | Percentage activity |  |  |
| --- | --- | --- | --- |
|  | 6 to 10 hours | 10 to 14 hours | 14 to 17.30 hours |
| Incubation | 21.5 | 25.1 | 27.0 |
| Perch on nest | 43.0 | 37.1 | 50.9 |
| Perch | 29.2 | 33.9 | 18.1 |
| Perch on ground | 0.8 | 1.7 | 1.8 |
| Feeding | 0.6 | 0.5 | 0.7 |
| Bath | 0.0 | 0.2 | 0.0 |
| Nest material collection | 0.4 | 0.3 | 0.0 |
| Nest arrangement | 2.9 | 0.7 | 0.8 |
| Mating | 1.6 | 0.5 | 0.7 |

**Figure 13.**
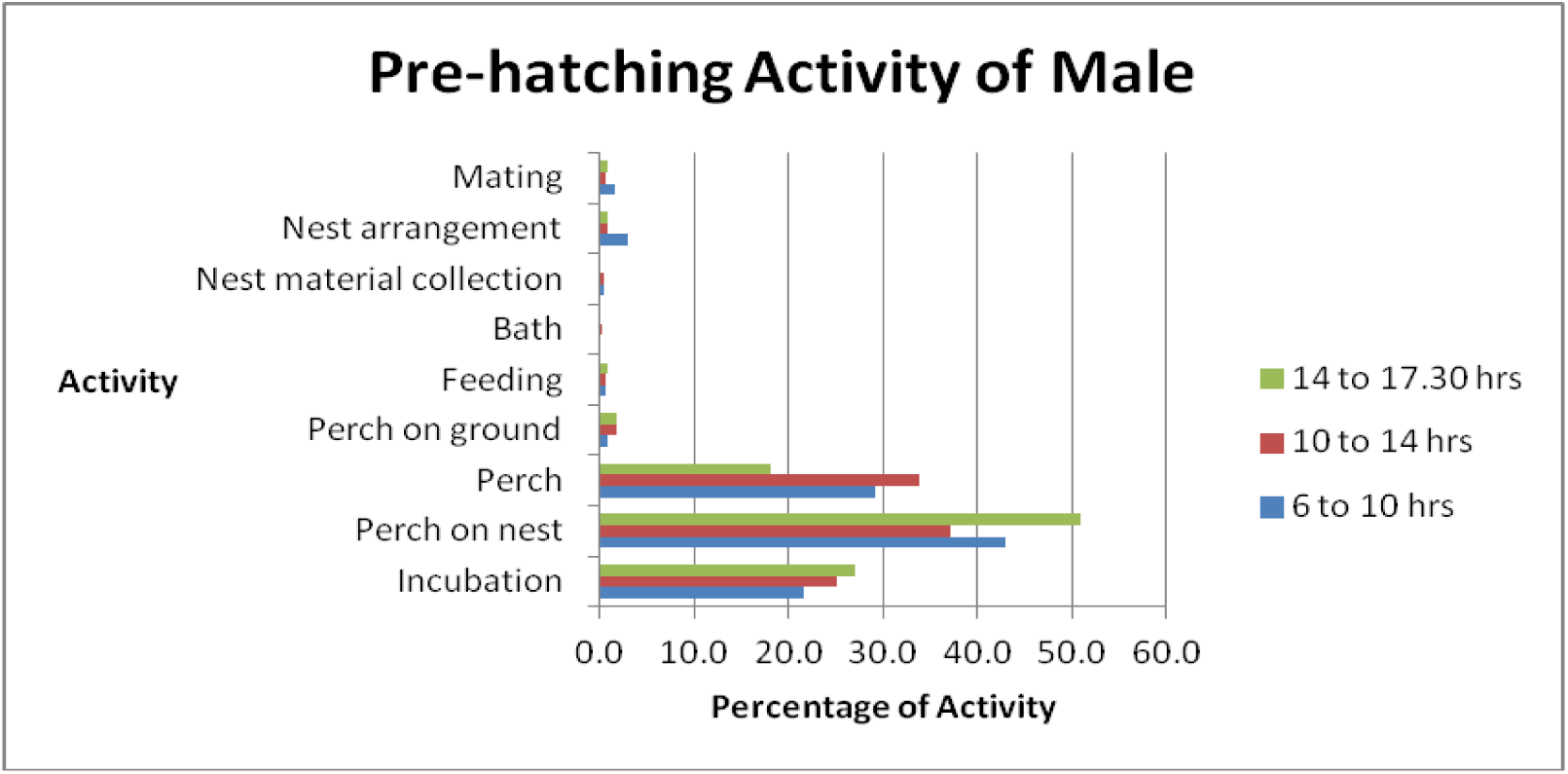
The activities of male Slender-billed vulture (Nest N7) during the pre-hatching period.

**Table 14.** The activities female Slender-billed vulture (Nest N7) during the pre-hatching period. Total observations of 20065 minutes.

| Activity | Percentage activity |  |  |
| --- | --- | --- | --- |
|  | 6 to 10 hours | 10 to 14 hours | 14 to 17.30 hours |
| Incubation | 15.1 | 37.2 | 30.8 |
| Perch on nest | 48.0 | 36.6 | 41.8 |
| Perch | 31.0 | 22.2 | 22.3 |
| Perch on ground | 0.2 | 1.6 | 1.0 |
| Feeding | 0.2 | 0.3 | 2.2 |
| Bath | 0.0 | 0.0 | 0.0 |
| Nest material collection | 0.1 | 0.1 | 0.2 |
| Nest arrangement | 3.7 | 1.5 | 1.0 |
| Mating | 1.7 | 0.5 | 0.7 |

**Figure 14.**
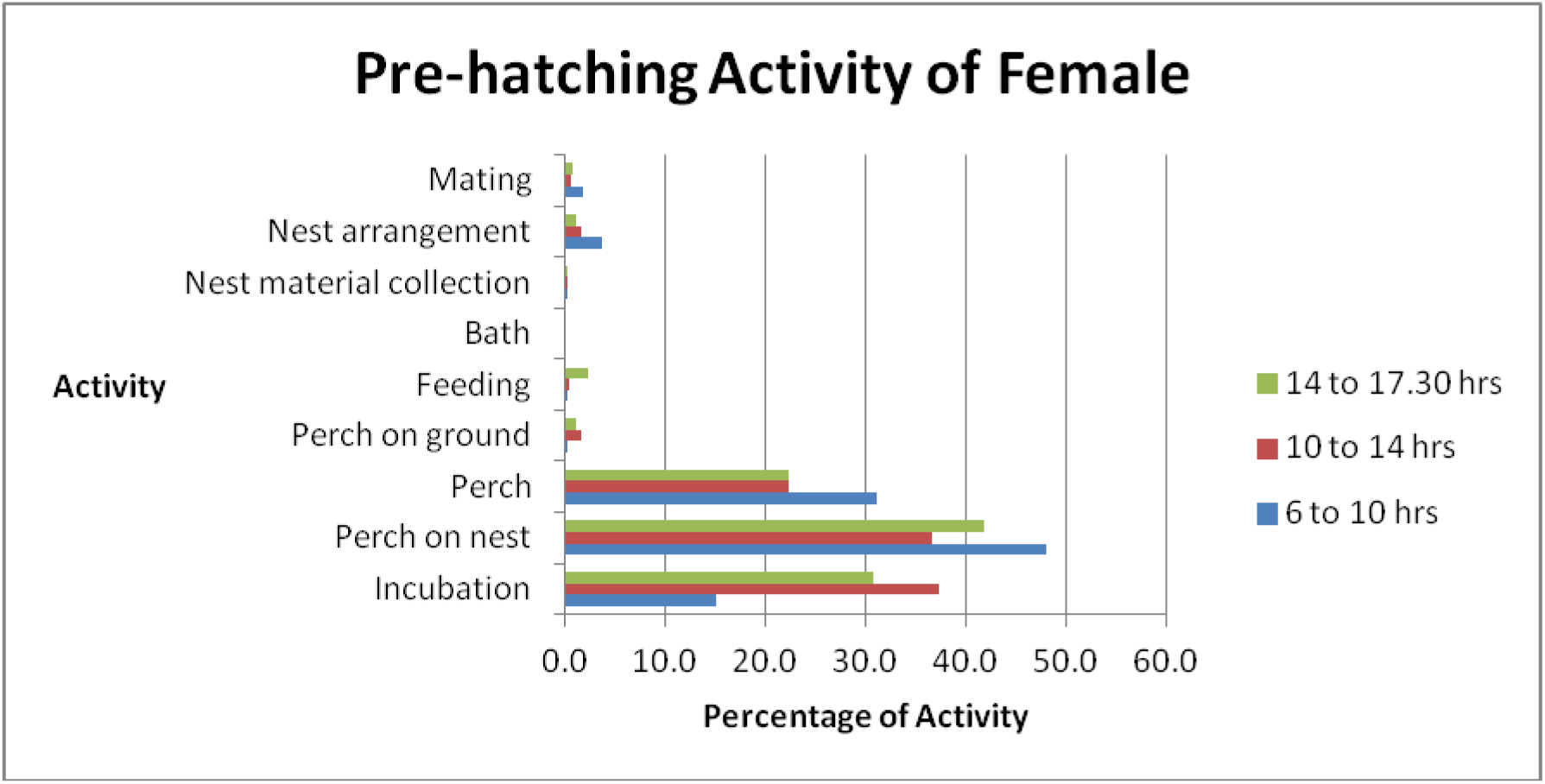
The activities female Slender-billed vulture (Nest N7) during the pre-hatching.

**Table 15.** The activities of male Slender-billed vulture (Nest N16) during the pre-hatching period. Total observations of 11035 minutes.

| Activity | Percentage activity |  |  |
| --- | --- | --- | --- |
|  | 6 to 10 hours | 10 to 14 hours | 14 to 17.30 hours |
| Incubation | 71 | 77.9 | 65.3 |
| Perch on nest | 5.7 | 4.5 | 6.1 |
| Perch | 19.1 | 11.8 | 21.7 |
| Perch on ground | 0.3 | 2.5 | 1.6 |
| Feeding | 1.3 | 1.3 | 3.8 |
| Bath | 0 | 0.1 | 0 |
| Nest material collection | 0.6 | 0.6 | 0.5 |
| Nest arrangement | 1.8 | 0.9 | 0.8 |
| Mating | 0.2 | 0.3 | 0.3 |

**Figure 15.**
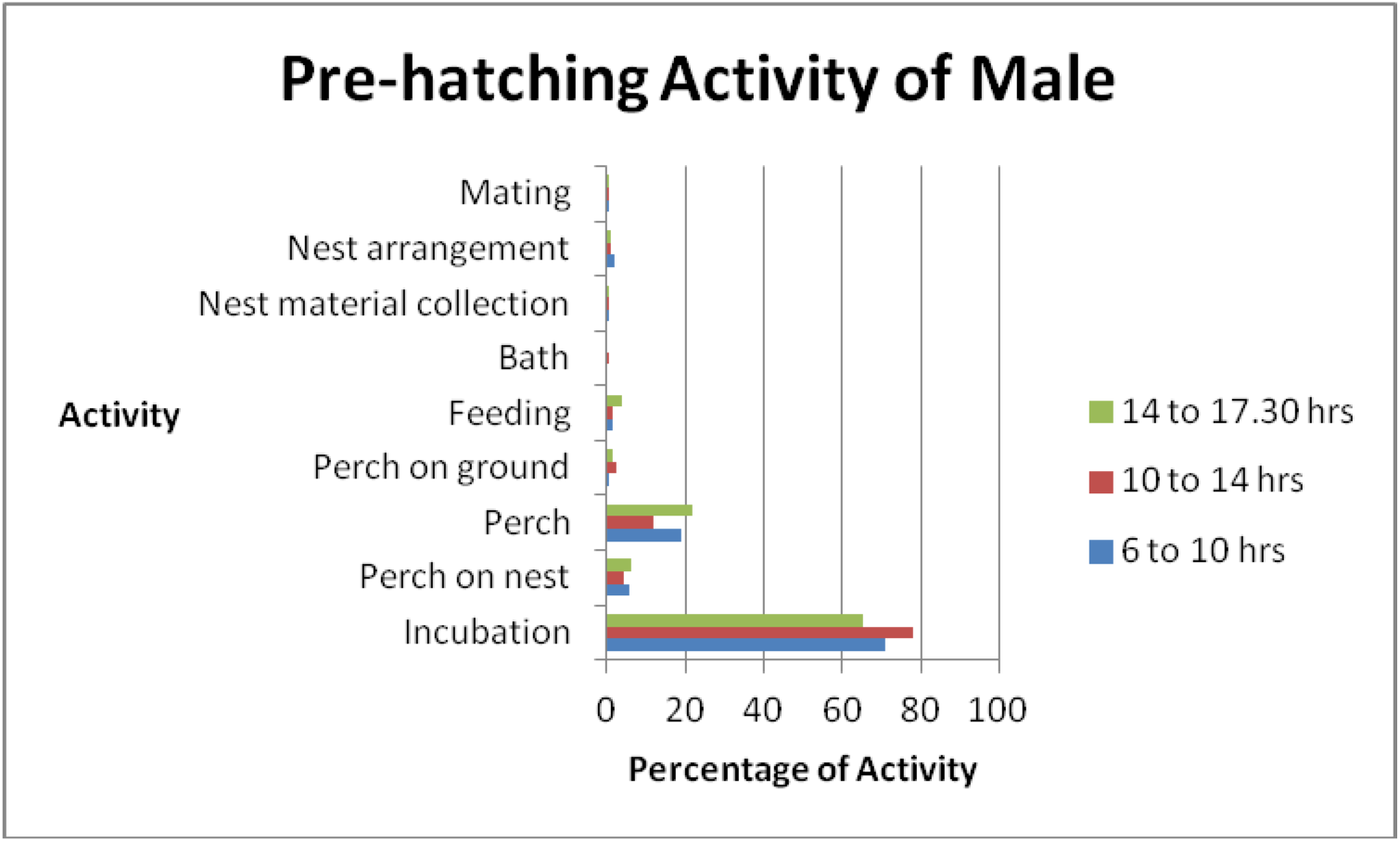
The activities of male Slender-billed vulture (Nest N16) during the pre-hatching.

**Table 16.** The activities of female Slender-billed vulture (Nest N16) during the pre-hatching period. Total observations of 8490 minutes.

| Activity | Percentage activity |  |  |
| --- | --- | --- | --- |
|  | 6 to 10 hours | 10 to 14 hours | 14 to 17.30 hours |
| Incubation | 63.1 | 58.4 | 49.4 |
| Perch on nest | 7.1 | 8.4 | 15.4 |
| Perch | 22.8 | 29.6 | 29.4 |
| Perch on ground | 0.7 | 1.4 | 0.7 |
| Feeding | 1.8 | 0.4 | 3.5 |
| Bath | 0 | 0 | 0 |
| Nest material collection | 0.2 | 0.3 | 0 |
| Nest arrangement | 4.2 | 1 | 1.1 |
| Mating | 0.2 | 0.4 | 0.4 |

**Figure 16.**
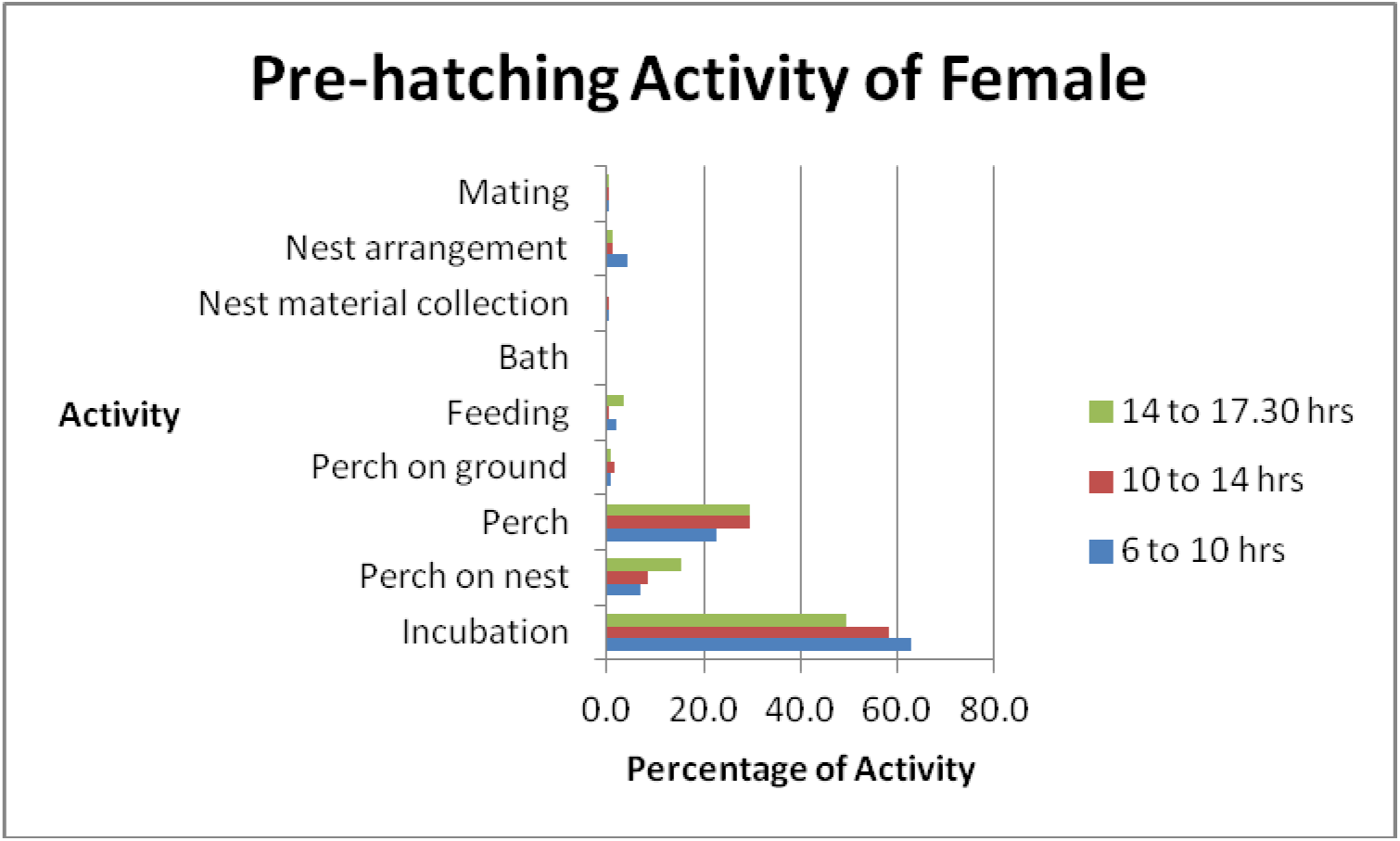
The activities of female Slender-billed vulture (Nest N16) during the pre-hatching period.

## Discussion

The study species included in this work, White-rumped vulture and Slender-billed vultures, were kept under captivity but under near natural condition with minimal human interference. All the observations were done through CCTV monitors from a distance so there was no change in normal behavior of the birds. Hence it was interesting to observe their time activity budget especially in breeding season. In the year 2009, a Slender-billed vulture pair and in 2010 a White-rumped vulture pair bred successfully at the VCBC for first time. In 2011, the three White-rumped and two Slender-billed vulture pairs were formed which laid eggs and were consistent in nest care throughout the breeding season.

The vultures were observed during day light hours for the study of their time budget and activity patterns. During the study of reintroduced California condors, the nests were observed during sunrise-sunset. In this case, the author considered that watching each nest for 40% of sunrise-sunset hours would provide sufficient observations of the subject (Sandhaus 2013). For studying behavior of the Greater spotted eagle, Graszynsky *et al* observed the raptor from 8 to 18 hours.

During this study the major activities of the breeding birds were incubation and brooding, followed by resting. For the nesting birds it was obvious to find major time spent on incubation and brooding when the eggs hatched, which are activities related to thermoregulation of the egg and nestling. Other than that, resting was observed as the most frequent activity and it stood second largest in the time budgets of breeding birds. Perching combined with the incubation and brooding activities emphasizes that most of the time spent by the adult birds is on nest. The other activities noted during breeding season such as feeding, nest building and maintenance and care of young occupied very small fraction of the time budget.

### Activities of adult vultures

#### 1. Incubation and brooding

During the breeding season incubation is the most important activity for growth of embryo inside the egg. It is well described regarding its definition, brood patch, physical and physiological factors, and role of parents (Rudolf Drent 1952). Proper incubation leads to successful hatching of the young one, which is an essential step in successful breeding. Brooding is an act of thermoregulation and protection for the hatchling and young nestling.

##### • Incubation and brooding in White-rumped vulture

In this study, the White-rumped vultures on an average spent 62.14% (n=6) of the total activity time for incubation. Male and female both spent almost similar time on incubation of the egg (61.72% in male and 62.5% in female). In all the time slots, the percentage of incubation remained fairly similar (morning (59.4%), noon (65.75%) and evening (61.2%)).

In White-rumped vulture, the overall average brooding time was 32% in male and 26% in female. The overall percentage of the brooding was noted fairly the similar throughout the day – 6-10 hours (33.8%), 10-14 hours (26.1%) and 14-17:30 hours (27.68%).

As the nestling grows, it develops feathers and manages to thermoregulate itself. This reduces its dependence on parent for brooding. In the late phase of nestling growth, the parents become less attentive to nest and nestling frequently perches away from the nest. Hence the average presence of parent on the nest during pre-hatching period (Incubation + Perch on the nest; 70.4%) is more than the average time spent during post-hatching period (Brooding + Perch on nest; 61.7%).

##### • Incubation and brooding in Slender-billed vulture

There were two pairs of Slender-billed vultures considered for the study. In both cases the egg didn’t hatch. The pair from nest 16 was successful in raising the nestling in earlier season. In this pair, male spend 71% and female spend 57% of the total activity time in incubation which was similar to that of White-rumped vulture pairs. Their incubation in three different periods of time during the day was fairly similar (6-10 hours (67%), 10-14 hours (68%) and 14-17:30 hours (57%)). The other pair of slender-billed from Nest 7 was inexperienced. The parents incubated the egg – male 24.5% and female 28%, spending most of the time perching on nest; probably causing the failure in hatching of the egg.

#### 2. Perch on nest

The vultures are well known to perch on tall trees at a vantage point. In captivity as well, they preferred higher perches and nest ledges during all seasons. In breeding season their perching on nest serves various purposes. It could be a short break from incubation or brooding to maintain right temperature of the egg or nestling, it could be aggression to intruders, and even could be stretching of legs and wings after a long span of incubation.

- During incubation, the White-rumped vultures perched on nest on an average for 8.27% of the total activity time observed. After hatching of the egg, the birds spent on an average 32.48% of the time perching on the nest. It could be the reason that the vultures required fewer efforts to keep the nesting warm as it was required during incubation, but perched on the nest guarding the nestling and nest. Depending on the atmospheric temperature, the vultures either sit tight or stand on the nest both while incubation or brooding.
- The Slender-billed vultures were observed to be perched on nest for 7.86% of the time.

#### 3. Resting (Perch)

Vultures spend considerable time in perching in the wild. In an aviary there are various kinds of perches made available at various heights for the vultures. The purpose of perching could be various – from active vigilance to rest. It was noticed that the perching is the second major activity in the captivity during the breeding season. A suitable perch provides sunlight to the vulture for sunning and also provides better view of the ground and air field. It could be indicator of hierarchy among vultures to gain higher and better perch.

- During incubation, the White-rumped vulture perched for on an average 19.87% while after hatching of egg, they were found perched 25.6% of the total time of observation in White-rumped vultures.
- The pairs of Slender-billed vultures observed to perch for 22.4% of the total time.

#### 4. Perch on ground

The activity of perch on ground or standing was analysed separately from perching on higher perches as they serve different purpose. The higher perches in nature serve for better and larger view of the area and security from ground dwelling predators, whereas perch on ground provides access to food and water. The ground also serves as a substrate to rest other than higher perches like trees and rocks. The vultures sit on ground for considerable time. It was noticed that after the birds in colony aviary settled, the White-rumped vultures often utilized the ground during sunning. A male vulture (B81) of nest 25was observed to sit/squat on ground whenever the female was busy in incubation (n= 30). The vultures also squat on ground when the dominant scavengers are feeding. Vultures were commonly sighted waiting for their turn, sitting on hocks, squatting and also sleeping on the ground while human were engaged in skinning the carcass of livestock or dogs/jackals/hyena were feeding on the carcass. Such incidences were observed at Golaghat, Goalpara and Dibrugarh districts of Assam (n=5).

- During this study, the breeding White-rumped vultures spent 4% of the time on ground during the incubation that reduced to 0.7% in post hatching period.
- The Slender-billed vultures spent an average of 1.19% time on the ground.

#### 5. Feeding

Although in nature, the vultures spend considerable time for foraging, the actual feeding takes really short time. In captivity, the food is available readily and assured. To mimic wild conditions, the food was provided on two days a week. It was observed that the breeding adults, especially during incubation and brooding spent less time on feeding. The feeding was carried out one by one by the breeding pair during incubation and brooding. It appeared that these birds were probably dominant and snatched food aggressively from the non-breeding and immature vultures. Similar observations are recorded in the mammalian scavenger-Spotted hyena *Crocuta crocuta* in Kenya where the lower ranking females spend more time in feeding than the high-ranking ones (Kolowski et al 2007). It has been noted that the herbivores spend more time of their activity for feeding. The time budget of herbivores such as in giraffe and elephants in Africa is well studied and shows feeding as the major activity. The generalist avian feeders also spend longer time on feeding as recorded in helmeted guinea fowl (*Numida meleagris*) (Kumasa & Bekele 2013). The vultures are adapted to feed only on dead animals. Vultures are also gregarious feeders. Although they compete with each other while feeding, they also depend on each other for the safety.

- The White-rumped vultures were seen feeding for on an average of 3.8% of the activity time during incubation. The breeding pair of vultures with nestling left the nest one by one to feed, spending an average of 0.6% of the total activity time for feeding.
- The Slender-billed vultures spent an average of 2% of the time for feeding where male and female spared almost equal time.

#### 6. Bath

Bathing is one of the maintenance activities. Bathing is very important in vultures as they feed on carcasses. There are chances of soiling the feathers while feeding and hence generally after the feeding the vultures take bath. It also helps them for to lower body temperature during summer. During this study the White-rumped vultures were seen spending 0.06% and Slender-billed vultures 0.02% of the activity time for bath. The bath time and period vary as per season and dominance of individuals. Yet more data would be required to reach any conclusion.

#### 7. Nest material collection

In the wild, the vultures collect twigs and small branches to construct the nest. For the captive vultures the nest material was provided on ground which was picked up by the breeding birds and carried to the ledges to construct nest. This was one of the important activities recorded during early breeding season. It was noticed that most of nest material was collected in early hours of the day and male individuals collected more nest material than the females. During the early phase of breeding i.e. in pre-hatching period, the male in White-rumped vulture was noted collecting nest material 3.5% of the time while females spent 0.5% (n=3) of time for the same. The Slender-billed vultures spent 2.4% of the time while the females spent only 0.9 % time for nest material collection (n=2). The activity was noticed in later months of breeding as well but with lower frequency.

#### 8. Nest material arrangement

Following the nest-material collection, both male and female arranged the material on the given ledges. The twigs were lifted in beak and were placed carefully. The birds sometimes sat to press the material to give desired shape to the nest. The nest construction was completed prior to the egg-laying. Once built, the nests needed to be maintained for holding the egg as well as the nestling till the nestling fledged out. In the White-rumped vulture, the nest material arrangement was carried out by female more than male (male 0.8%, female 1.19% and an average of 1% of the total time). This activity remained consistent at 1% during pre and post hatching periods.

In case of Slender-billed vultures, the nest material arrangement was noticed for 1.63% of the total time and female arranged it for more time (1.25%) than male (0.8%).

#### 9. Coition

Coition is a short time activity, which is sometimes followed by preening. Mating is an essential act in breeding as well as for bond formation and maintenance between the breeding pair. Mating events were observed more during the breeding season but it continued even in non-breeding season. Mating was observed in the breeding birds, even on the day of hatching of the egg, when their nestling was grown up and was in nest, and also after their nestling fledged.

- In White-rumped vulture it was noted that mating accounted for 0.5% of the time during incubation that reduced to 0.1% during the post hatching period.
- In Slender-billed vultures, mating was recorded as 0.3% of the total time.

#### 10. Feeding the nestling

In the post hatching period, feeding the nestling is the main activity on the nest of the parents. Both male and female take equal care of the nestling and feed it. In addition to the physical development of the young bird, these activities develop a bond between the young one and its parents. The White-rumped vultures were seen feeding their nestling for 7.84% of the time.

#### 11. Preening of the nestling

Other than feeding, preening is an essential activity of parent birds to establish bonding between parent and offspring. This activity by the parents controls the ecto-parasites on the hatchling’s body, which otherwise the nestlings cannot take care on its own. Preening maintains the shape and cleanliness of feathers and the young ones need to learn this act which they would practice for life-long. In the White-rumped vultures, the parent birds were seen preening for 0.4% of the time.

### Activities of Nestlings

The focus of this study was on the breeding birds, yet, the activities of nestlings were recorded opportunistically in the White-rumped vulture’s nests. The difficulties in observing them were – the hatchling and early-stage nestlings are too small to observe in a nest, even with a CCTV camera in the captivity. Most of the time they are hidden under the wings of brooding birds and their activities could not be observed directly. During the four to five months in the nest, from hatching until fledgling, the nestling went through various phases of growth. The young one’s alertness and activities changed from time to time. On the whole, the nestlings rested most of time which accounts to an average of 63% of the total time of observation. Nestlings remained perched on the nest for about 23.75% of the time. They were fed by parent birds for about 4.6% of the time while 2% of time they were recorded calling and begging for food.

## Conclusion

The study of time budget of White-rumped and Slender-billed vultures during breeding season revealed that the birds spend major time in incubating and brooding at the nests. Both sexes incubated and brood for similar time periods. Their attentiveness to nest was seen throughout the observation hours of day. During the breeding season, the nest ledge was their preferred perch. They used other perches and ground as well for perching but it was only for a short duration. Nest material collection was carried out by both sexes but male contributed slightly more. The nest arrangement was carried out by both parents during incubation as well as during raising the nestling. It appeared that the other activities like feeding, bath were well organized giving prime importance to the care of nest and the nestling.

The time activity budget studies confirmed the equal role of male and female for sharing the duties of incubation and brooding in White-rumped vulture and Slender-billed vulture. It unfolded some interesting facts such as the males give more efforts and time to collect the nest material while the females spend more time to arrange it. As observed in wild, in captivity also, one of the parents was noted taking care of the egg or nestling while the other perching at nearby perch or even on the ground. These observations suggests that the vultures maintained for breeding purpose had their instincts and behavior similar to the wild birds in the nature, and the data could be used as baseline for the future studies.

